# BellaVista: Open-Source Visualization for Imaging-Based Spatial Transcriptomics

**DOI:** 10.1101/2025.01.07.631783

**Authors:** Annabelle M. Coles, Yuening Liu, Pallav Kosuri

## Abstract

Imaging-based spatial transcriptomics can reveal gene expression *in situ* by locating and identifying individual RNA molecules at subcellular resolution. These datasets typically contain an abundance of information that when analyzed appropriately can reveal tissue organization across scales from molecules to entire organs. However, there is currently a lack of simple open-source tools that facilitate visualization, quality control, and custom spatial analysis.

Here we introduce BellaVista, a lightweight open-source tool for interactive visualization and exploration of imaging-based spatial transcriptomics data. BellaVista natively supports data from Xenium (10x Genomics), MERSCOPE (Vizgen), and custom MERFISH platforms. By providing a simple means for simultaneous visualization of images, transcripts, and cell segmentation boundaries, we anticipate that BellaVista will accelerate accessibility, exploration and discovery in the rapidly expanding field of spatial biology. BellaVista is available at https://github.com/pkosurilab/BellaVista.

## 1 Introduction

Imaging-based spatial transcriptomics makes it possible to measure gene expression across space in biological samples by locating and identifying single RNA molecules (Tian, et al., 2023). Initially introduced in cell culture (Chen, et al., 2015; Lubeck, et al., 2014), these methods have more recently been used to reveal spatial patterns of gene expression in sections of intact tissue such as brain (Eng, et al., 2019; Moffitt, et al., 2018; Zhang, et al., 2023), heart (Farah, et al., 2024), intestine (Cadinu, et al., 2024; Reina-Campos, et al., 2024), liver (Watson, et al., 2024), and cancer tumors (Chen, et al., 2024). These studies have already led to many discoveries including fundamental principles of functional organization in tissue, spatial niches that drive organ development, and intratumoral regions associated with favorable outcome of immunotherapy. Furthermore, thanks to the public accessibility of the raw data from these studies, many more discoveries will likely be made as more researchers are able to access and analyze these often vast and only partially explored datasets. Crucially, initial as well as subsequent analysis of image-based spatial transcriptomics data depend on user-friendly software that enables researchers to visually inspect multi-modal data, perform quality control, and generate and test physiologically motivated hypotheses.

Current visualization tools designed for interactive exploration of single-cell spatial transcriptomic data include SpatialDB (Fan, et al., 2019), Samui (Sriworarat, et al., 2023), SpatialData (Marconato, et al., 2024), and Vitessce (Keller, et al., 2024). Of these, SpatialDB and Samui plot spatial gene expression at a cellular level but currently do not support simultaneous visualization of multiple genes, individual transcript locations, or cellular segmentations. Vitessce and SpatialData can additionally be used to plot transcript locations and cell segmentations; however, these tools aggregate transcript data for all genes and plot them in a single layer. Thus, they cannot be used to readily visualize transcript locations solely for a single gene or a subset of genes, making it challenging to identify gene-specific spatial expression patterns. Commercial spatial transcriptomic platforms such as Xenium (10x Genomics) and MERSCOPE (Vizgen) feature their own proprietary software, however, the closed-sourced nature of these tools limit their utility for the research community. Consequently, there is a lack of open-source tools that both utilize the comprehensive information of a spatial transcriptomic dataset and support native spatial transcriptomic data file formats across platforms. To address this need, we developed BellaVista, a lightweight open-source tool for interactive data exploration of imaging-based spatial transcriptomics data. BellaVista can be used for many essential analysis tasks such as identifying spatial gene expression patterns, evaluating cell segmentation accuracy, and validating spatial analysis discoveries.

## 2 Visualization framework

BellaVista is a command-line Python application that utilizes the napari GUI (Sofroniew, 2024) to enable interactive visualization of multimodal spatial transcriptomics data including images, gene transcript locations, and cell and/or nuclear segmentations. Image, gene, and segmentation data are displayed as separate layers, with individual layers for each gene. Within the napari GUI, a user can pan and zoom in/out to explore large and complex multimodal data. Furthermore, a user can toggle layers on and off to visualize and identify different spatial patterns of gene expression. Importantly, via napari, BellaVista can be used to create and reproduce publication-quality figures and animations. Additional usage details and tutorials are provided in the user guide accompanying this manuscript (Supplementary File S1).

## 3 Methods

### 3.1 Installation, configuration and usage instructions

BellaVista is available as a Python package and can be installed on Windows, macOS, and Linux via the package manager pip. BellaVista is run from the command line and accepts a JSON configuration file that specifies the path to the input data, and parameters for visualization. Images, gene transcript locations, and segmentations are accepted as input data. If a feature is not provided, BellaVista will skip the processing and visualization of these data. Documentation, tutorials, and example JSON configuration files are available on GitHub at https://github.com/pkosurilab/BellaVista.

### 3.2 Input data

BellaVista natively supports spatial transcriptomics data from Xenium (10x Genomics), MERSCOPE (Vizgen), and custom MERFISH platforms. The platform used and paths to the required input files can be specified by the user in a JSON configuration file. Regardless of which platform was used to acquire the data, BellaVista will generate a standardized dataset in a simple format that is used for visualization. This standardized version of the data is generated only once, when loading the input data for the first time. The standardized data are stored locally and then accessed for rapid loading in subsequent sessions.

To generate the standardized BellaVista data format, image files are converted to a pyramidal OME-Zarr next-generation file format (NGFF) utilizing compression of image chunks at multiple resolution levels for rapid loading and smooth transitioning between different magnifications as the user zooms in and out (Moore, et al., 2023). Gene transcript locations are stored in a dictionary mapping transcript coordinate locations to gene identities, and segmentations are stored in an array. These standardized data are saved to a BellaVista subfolder within the input data directory. Typically, the standardized dataset is equal to or smaller in size than the input dataset; therefore, we suggest having available disk storage equivalent to the size of the input data. Additionally, when loading a typical-sized dataset, at least 16 GB of RAM is recommended.

## 4 Use case

To demonstrate how BellaVista can be used to visualize imaging-based spatial transcriptomics data, we explored Xenium data of the mouse brain (https://doi.org/10.5281/zenodo.14279832). A JSON configuration file detailing the paths to the Xenium input data and selected visualization parameters was passed to BellaVista. After the output data had been generated, image and transcript data were displayed at a zoomed-out magnification (Fig. 1B). At this low level of magnification, it was possible to see prominent spatial patterns of gene expression across the tissue. To investigate the data at the scale of single cells, we then zoomed in to a higher magnification and visually inspected each modality in the dataset. By toggling on only the image layer, cell locations became visible in the morphology image showing DAPI-stained nuclei (Fig. 1C). We then switched on the cell boundaries layer, and used the boundary and DAPI overlay to evaluate the accuracy of cell segmentation (Fig. 1D). A single nucleus was visible within each cell boundary, suggesting a baseline accuracy of the segmentation data. Finally, toggling on the transcript layers revealed subcellular spatial gene expression patterns, with some transcripts appearing enriched in the nuclei while others appearing evenly distributed within the cells (Fig. 1E). With the ability to easily visualize salient features at various magnifications, BellaVista provides an effective platform for investigating imaging-based spatial transcriptomics data.

**Figure 1.**
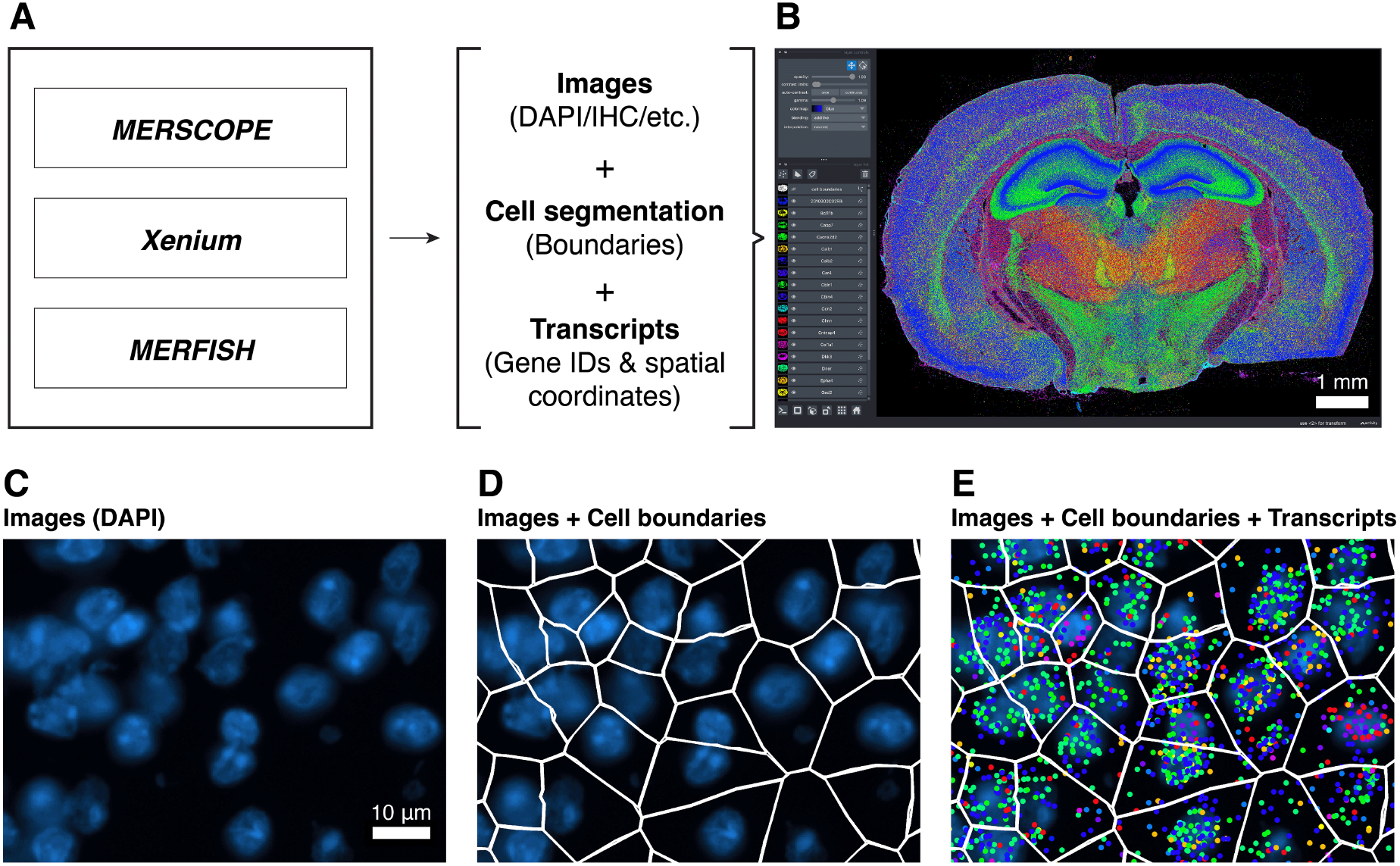
BellaVista enables visualization of imaging-based spatial transcriptomics data. (A) Datasets can be imported from any of the main platforms for imaging-based spatial transcriptomics, including MERSCOPE (Vizgen), Xenium (10x Genomics), and custom MERFISH platforms. BellaVista imports images, cell segmentation boundary data, and gene transcript localization data. Using these inputs, a standardized dataset is created that enables rapid, platform-agnostic visualization in napari. (B) BellaVista rendering of Xenium data from a coronal section of mouse brain, revealing tissue-wide spatial gene expression patterns. The visualizer can be used to browse image data (C), validate cell segmentation boundaries (D), and inspect single-cell transcription profiles (E).

## 5 Summary

BellaVista is a lightweight open-source visualizer for imaging-based spatial transcriptomics data, currently compatible with the Xenium, MERSCOPE, and custom MERFISH platforms. The software can be used to explore large multimodal datasets and discover patterns across modalities including imaging data, transcript locations, and cell segmentation boundaries. We have found it to be particularly useful for evaluating the accuracy of cell segmentation, for identifying subcellular patterns of transcript distributions, and to discover non-random gene expression patterns. Through integration with napari, users can make publication-quality figures, generate animations, and add their own plugins for custom analysis. Single-cell spatial transcriptomics is a rapidly growing field that is set to transform how we investigate complex biological systems. With the increase in available spatial data, we foresee that BellaVista will become increasingly useful in the pursuit of new insights into fundamental biology and human health.

## Supplementary data

Supplementary data consist of a user manual and are provided along with this manuscript.

## Acknowledgements

We would like to thank the members of the Kosuri Lab who tested BellaVista and provided feedback.

## Data availability

The Xenium (10x Genomics) mouse brain dataset used for the figures is available on Zenodo: https://doi.org/10.5281/zenodo.14279832

