## Supplementary File S1 for "BellaVista: Open-Source Visualization for Imaging-Based Spatial Transcriptomics"

KOSURI
LAB

### BellaVista

#### User Guide

Version 0.1

Prepared by
Annabelle M. Coles

### 1 Application Overview

BellaVista is a command-line application written in Python, utilizing the napari GUI [1] for interactive visualization of multimodal spatial transcriptomic data including images, gene transcript locations, and cell/nuclear segmentations. Image, transcript, and segmentation data are displayed as separate layers, with individual layers for each gene. Within the napari GUI, a user can pan and zoom in/out to explore large and complex multimodal data. Furthermore, a user can toggle layers on and off to visualize and identify different spatial patterns of gene expression. Importantly, via napari, BellaVista can be used to create and reproduce publication-quality figures and animations. Installation, usage, and figure generation instructions can be found in subsequent chapters.

Currently, BellaVista accepts spatial transcriptomics data from 10x Genomics Xenium, Vizgen MERSCOPE, and custom MERFISH platforms. As the BellaVista package is compatible with multiple platforms, users utilizing data from various platforms can use BellaVista to easily explore all datasets within a single application.

The BellaVista package was designed as a lightweight application, making it easier for users to customize the code for custom setups, analyses, and needs. Check out the documentation website for API details, and the napari website for further documentation and features you can implement!

**Links**

BellaVista GitHub: https://github.com/pkosurilab/BellaVista
BellaVista documentation website: https://bellavista.readthedocs.io
napari: https://napari.org/

*Figure 1 on next page ⟶*

**Figure 1: BellaVista enables visualization of imaging-based spatial transcriptomics data**

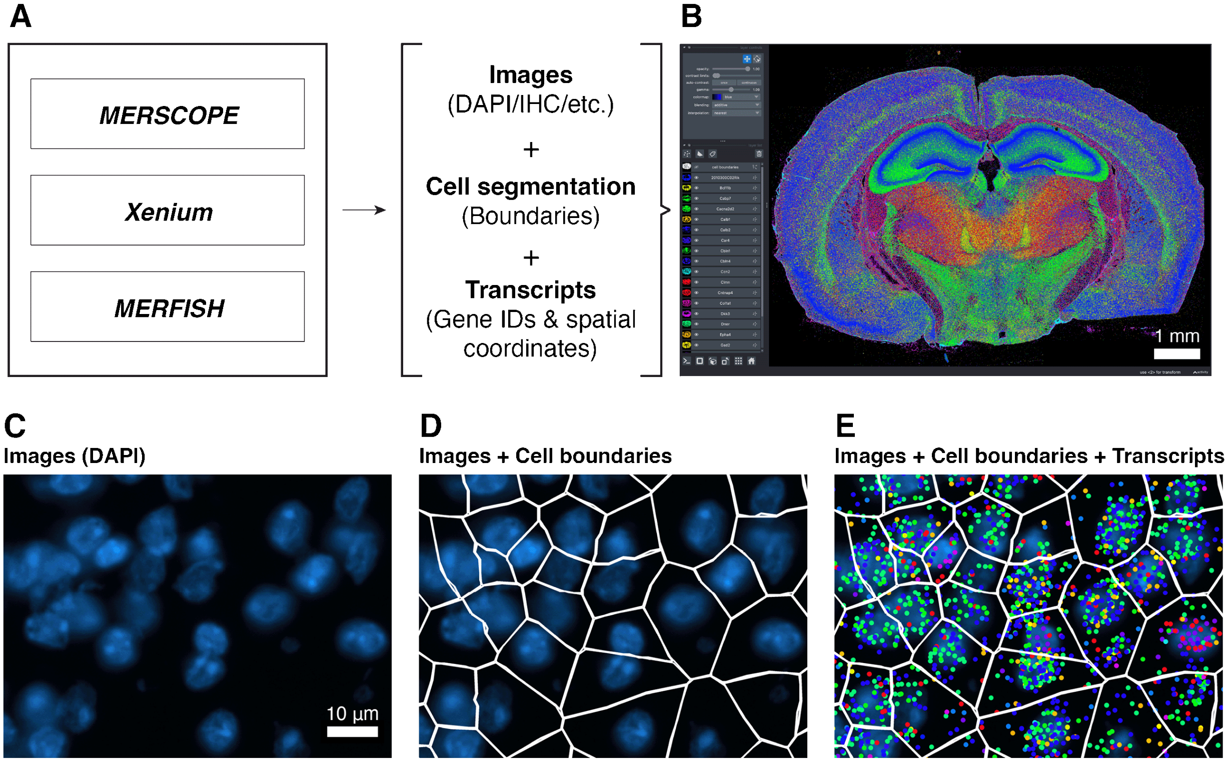

(A) Datasets can be imported from any of the main platforms for imaging-based spatial transcriptomics, including MERSCOPE (Vizgen), Xenium (10x Genomics), and custom MERFISH platforms. BellaVista imports images, cell segmentation boundary data, and gene transcript localization data. Using these inputs, a standardized data format is created that enables rapid, platform-agnostic visualization in napari. (B) BellaVista rendering of Xenium data from a coronal section of mouse brain, revealing spatial patterns of gene expression. The visualizer can be used to browse image data (C), validate cell segmentation boundaries (D), and inspect single-cell transcription profiles (E).

### 2 Installation

The following instructions require that you have Anaconda installed. It is recommended to create an Anaconda virtual environment to prevent conflicting package dependencies. The package can be installed from PyPI via pip (recommended) or from the GitHub repository. BellaVista requires Python 3.9 or above and is dependent on GPU for rendering.

In the terminal, create and activate a new virtual environment:

```
conda create -n bellavista_env python=3.12
conda activate bellavista_env
```

Installation via pip:

```
pip install bellavista
```

Alternatively, you can clone the repository from GitHub:

```
conda install git
git clone https://github.com/pkosurilab/BellaVista
pip install -e BellaVista
```

Or, you can download and install using the source code from GitHub:

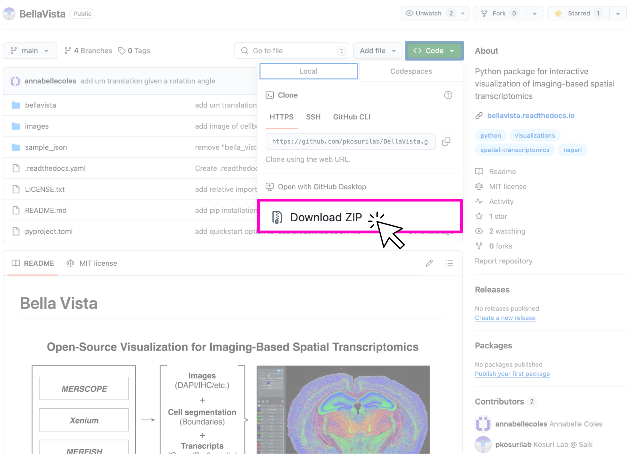

**Tip**: using the `pip install -e` flag will allow you to edit the source code!

Anaconda download: https://www.anaconda.com/download
GitHub: https://github.com/pkosurilab/BellaVista
PyPI package: https://pypi.org/project/bellavista

#### 2.1 Quick Start

Refer to this short tutorial for loading BellaVista with a sample Xenium dataset. This dataset includes DAPI morphology images, cell/nuclear segmentations, and transcript locations collected from the mouse brain.

1. Download and unzip the dataset folder "xenium_mouse_brain_rep3.zip" from Zenodo. Link to dataset: https://zenodo.org/records/14279832

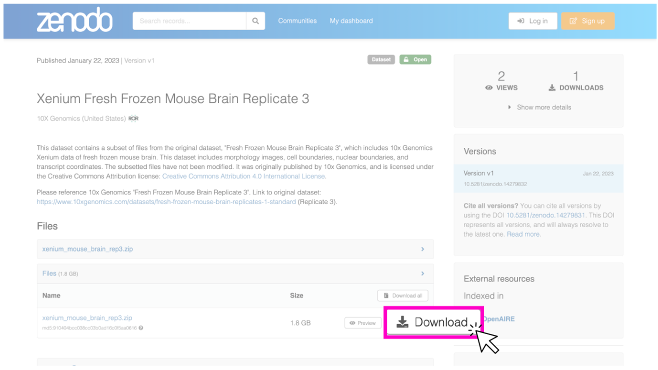

2. In the terminal, run BellaVista using the following command, replacing "/path/to/xenium_mouse_brain_rep3" with your local path to the downloaded, unzipped, sample dataset folder.

```
bellavista --xenium-sample /path/to/xenium_mouse_brain_rep3
```

**Note**: It will take a few minutes to create the required data files. The terminal will print updates & have progress bars for time consuming steps.

3. Once successfully loaded, you should see the message `Data Loaded!` in the terminal. A napari window should appear displaying the data similar to the image below:

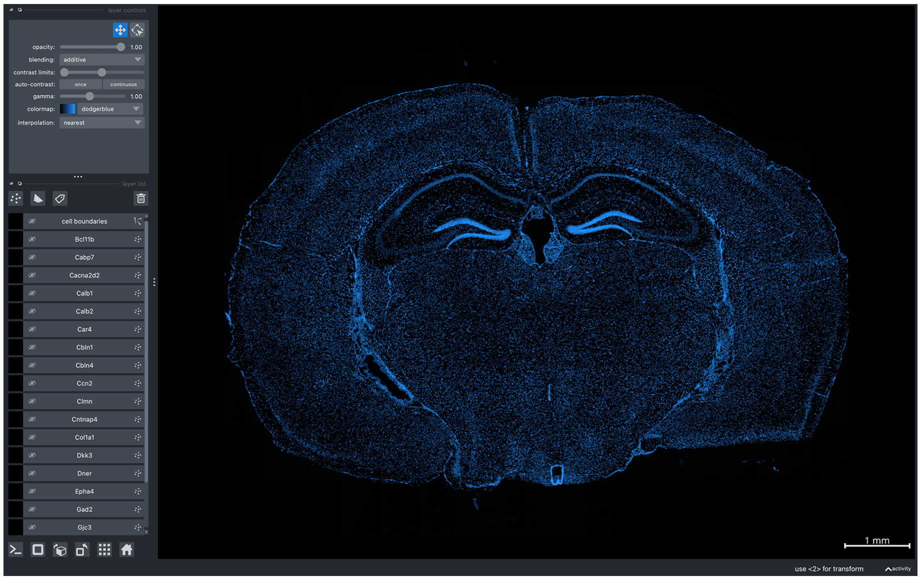

**Tip**: This is a large dataset, so if the program encounters a memory-related error, try visualizing a smaller subset of the data:

```
bellavista --xenium-sample-lite /path/to/xenium_mouse_brain_rep3
```

Using the mouse/trackpad to zoom in & out; click & drag to move around the canvas, you can interactively move around the napari canvas to explore the data. Try zooming in & out, toggling layers on & off to see different spatial features!

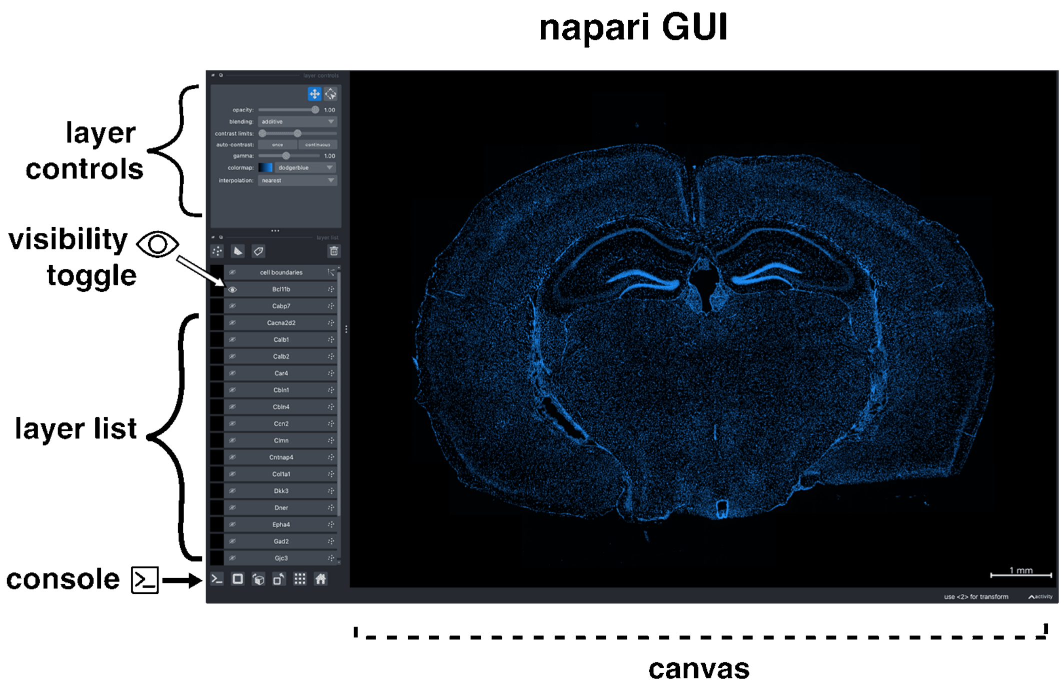

**Figure 2:**

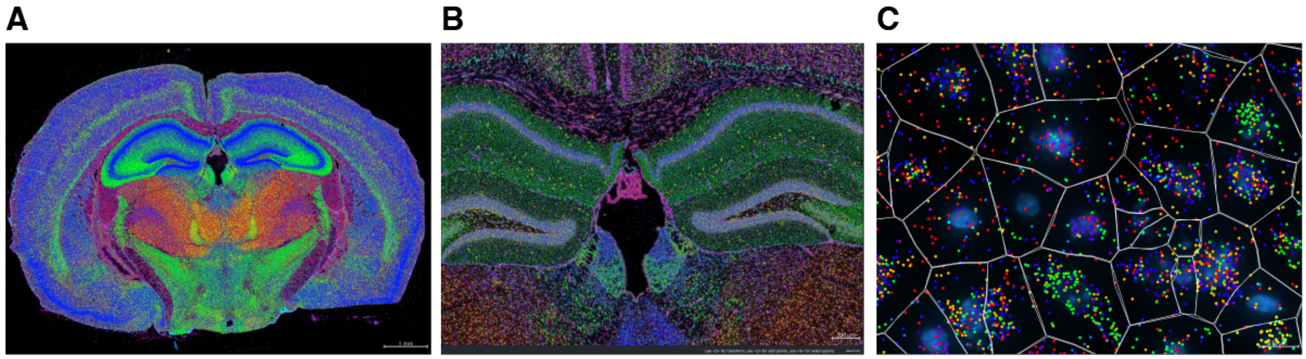

(A) Selected gene layers are toggled on to highlight spatial gene expression patterns.
(B) Zoomed-in image of spatial gene expression patterns. (C) Further zoomed-in, higher magnification screenshot of DAPI image, transcripts, and cell segmentations.

### 3 JSON Configuration

BellaVista uses a dataset-specific JSON configuration file to generate unified data formats and visualize the dataset. The JSON configuration file has three sections of parameters (1) General parameters, (2) Visualization parameters, and (3) Input file parameters. Below are descriptions and accepted values for each parameter; parameters with a defined *default* value are optional.

**JSON notes**: Boolean values are lowercase, and should be written as: `true/false` not `True/False`. Strings must be wrapped in double quotation marks, not single quotes, i.e. the correct format is: `"xenium"` not `'xenium'`

#### 3.1 General Parameters

These general parameters are used to indicate where to access and what platform was used to obtain the data. These are *required* for generating the output formats needed for visualization.

---

**system:** *string*

Specifies the spatial transcriptomic technology/platform. Allowed values: "Xenium", "MERSCOPE", or "MERlin" (for custom MERFISH).

- *The input is not case-sensitive, so values "xenium", "Xenium", and "XENIUM" are treated equivalently.*

**data_folder:** *string*

The path to the folder where the dataset data files are stored. BellaVista visualization files will be saved in a new subfolder "BellaVista_output" within data_folder.

**create_bellavista_inputs:** *boolean, default=true*

Specifies whether to generate the necessary visualization files for BellaVista. It should be set to `true` when loading the data for the first time. It can be set to `false` in later runs, as the files will already have been created.

- *If set to `true` and the visualization files already exist from a previous run, BellaVista will skip recreating those files and only generate any missing ones.*

#### 3.2 Visualization Parameters

Optional visualization parameters can be defined to customize how the data is loaded and displayed. The parameters below determine if a feature is loaded into the current viewing session. Reducing the number of features visualized will reduce the amount of computer memory required, and consequently reduce loading times and improve navigation performance. Additionally, a subsetted list of genes can be defined, and only data for these genes will be loaded into the current viewing session.

---

**plot_image:** *boolean, default=false*
Display image(s).

**plot_transcripts:** *boolean, default=false*
Plot spatial coordinates of gene transcripts.

**plot_allgenes:** *boolean, default=true*
Plot transcripts for all gene IDs. If set to `false`, only the gene IDs specified in selected_genes will be plotted.

**genes_visible_on_startup:** *boolean, default=false*
Controls the visibility of all gene layers at startup. If set to `false`, the gene layers will be hidden.

- *Setting this option to `false` improves navigation performance. Gene layers can be shown later using the visibility toggle button.*

**selected_genes:** *1D array of strings, default=None*
Specifies the gene IDs whose transcripts will be plotted. If not provided, transcripts for all genes will be plotted.

**plot_cell_seg:** *boolean, default=false*
Plot cell segmentation boundaries.

**plot_nuclear_seg:** *boolean, default=false*
Plot nuclear segmentation boundaries.

*Continued on next page ⟶*

Other visualization parameters are used to customize how the data is displayed. These parameters can be easily and dynamically changed during a viewing session in the napari GUI and console.

---

**image_colormap:** *1D array of strings, default=None*
Specifies the image colormaps to be used for the image(s). The colormaps will be applied to the corresponding image, at the same list index/position, as the list of images given in the input file parameters.

- *If there are multiple images, but only a single value is given, the specified colormap will be used for all images.*

**transcript_point_size:** *float, default=1*
Size of the points representing individual transcript coordinates.

**contrast_limits:** *tuple array of integers, default=None*
Range of values [0, 65535] used to set the contrast limits for the displayed image(s).

**rotate_angle:** *integer, default=0*
Rotation angle in degrees, within the range [0, 360], by which to rotate the data.

#### 3.3 Input File Parameters

The input file parameters indicate which data should be processed and subsequently visualized. When processing each feature, if a required input file is missing or an error occurs, BellaVista will automatically skip the processing and visualization of this feature. More information about data processing steps and outputs can be found in section (3.4) Output Formats.

As each platform organizes their datasets differently, the platforms have different platform-specific input file requirements. Please refer to the relevant sections for your platform(s).

| Platform | Pages |
|---|---|
| Xenium | 12 – 13 |
| MERSCOPE | 14 – 15 |
| Custom MERFISH | 16 – 19 |

##### 3.3.1 Xenium

Below is a description of input file parameters for visualizing 10x Genomics Xenium data. All input parameters are optional; if you are missing some input files, remove those input file parameters from the JSON file. BellaVista will skip the visualization of these features.

**Important:** All input file paths must be relative paths to *data_folder*.

- *data_folder* = the path to the folder where the dataset data files are stored.

---

**transcript_filename:** *string, default=None*
Relative path to a Parquet or CSV file containing transcript spatial locations. If no file is provided, transcripts will not be processed or visualized.

**images:** *string or 1D array of strings, default=None*
Relative path to image file(s). Must be an OME-TIFF or TIFF file. If no file is provided, images will not be processed or visualized.

- *When visualizing a single image, provide the file path as a string. For multiple images, pass them as a list of filenames. For example, use "DAPI.tif" for a single image or ["DAPI.tif", "PolyT.tif"] for multiple images.*

**z_plane:** *integer, default=0*
Z-plane of image to be visualize. If no value is provided, the first z-plane will be used.

**cell_segmentation:** *string, default=None*
Relative path to Parquet or Zarr file containing cell segmentations. If no file is provided, cell segmentations will not be processed or visualized.

**nuclear_segmentation:** *string, default=None*
relative path to Parquet or Zarr file containing nuclear segmentations. If no file is provided, nuclear segmentations will not be processed or visualized.

*Continued on next page ⟶*

Example Xenium JSON configuration file:

```
{
    "system": "xenium",
    "data_folder": "/path/to/xenium_brain_rep3",
    "create_bellavista_inputs": true,

    "visualization_parameters": {
        "plot_image": true,
        "plot_transcripts": true,
        "plot_cell_seg": true,
        "contrast_limits": [0, 5000],
        "rotate_angle": 180
    },

    "input_files": {
        "transcript_filename": "transcripts.parquet",
        "images": "morphology_mip.ome.tif",
        "cell_segmentation": "cell_boundaries.parquet"
    }
}
```

Additional Xenium JSON configuration files and tutorials can be found on the documentation website: https://bellavista.readthedocs.io

#### Next steps:

Once your JSON is correctly configured for your dataset, you can run BellaVista in the terminal:

```
bellavista my_dataset.json
```

- *Replace "my_dataset.json" with the filename of the JSON you created. The JSON file argument should contain the file path to your JSON file.*

Skip to section **(4) Running BellaVista** for more information!

##### 3.3.2 MERSCOPE

Below is a description of input file parameters for visualizing Vizgen MERSCOPE data. All input parameters are optional; if you are missing some input files, remove those input file parameters from the JSON file. BellaVista will skip the visualization of these features.

**Important:** All input file paths must be relative paths to *data_folder*.

- *data_folder* = the path to the folder where the dataset data files are stored.

---

**transcript_filename:** *string, default=None*
Relative path to CSV file containing transcript spatial locations. If no file is provided, transcripts will not be processed or visualized.

**images:** *string or 1D array of strings, default=None*
Relative path to image file(s). Must be a TIFF file. If no file is provided, images will not be processed or visualized.

- *When visualizing a single image, provide the file path as a string. For multiple images, pass them as a list of filenames. For example, use "DAPI.tif" for a single image or ["DAPI.tif", "PolyT.tif"] for multiple images.*

**um_to_px_transform:** *string, default=None*
Relative path to CSV file containing micron to pixel transformations. This is *required* for displaying images. If no file is provided, no images can be visualized.

**z_plane:** *integer, default=0*
Z-plane of segmentations to plot. We suggest this match the z-plane of the image you are visualizing. If no value is provided, the segmentations from the first z-plane will be processed and visualized.

**cell_segmentation:** *string, default=None*
Relative path to cell segmentations. This must be a Parquet file (MERSCOPE v232 or later) or a folder containing HDF5 files (pre v232). If no file/folder path is provided, cell segmentations will not be processed or visualized.

**nuclear_segmentation:** *string, default=None*
Relative path to nuclear segmentations. This must be a Parquet file (MERSCOPE v232 or later) or a folder containing HDF5 files (pre v232). If no file/folder path is provided, nuclear segmentations will not be processed or visualized.

*Continued on next page ⟶*

Certain features require multiple input files. Summary of required input files for each feature:

**Transcripts**: "transcript_filename"

**Images**: (1) "images" **&** (2) "um_to_px_transform"

**Segmentations**: "cell_segmentation" and/or "nuclear_segmentation"

Example MERSCOPE JSON configuration file:

```
{
    "system": "merscope",
    "data_folder": "/path/to/vizgen_brain_s1r1",
    "create_bellavista_inputs": true,

    "visualization_parameters": {
        "plot_image": true,
        "plot_transcripts": true,
        "plot_cell_seg": true
    },

    "input_files": {
        "transcript_filename": "detected_transcripts_S1R1.csv",
        "images":  "mosaic_DAPI_z3.tif",
        "um_to_px_transform": "micron_to_mosaic_pixel_transform.csv",
        "z_plane": 3,
        "cell_segmentation": "cell_boundaries"
    }
}
```

Additional MERSCOPE JSON configuration files and tutorials can be found on the documentation website: https://bellavista.readthedocs.io

#### Next steps:

Once your JSON is correctly configured for your dataset, you can run BellaVista in the terminal:

```
bellavista my_dataset.json
```

- *Replace "my_dataset.json" with the filename of the JSON you created. The JSON file argument should contain the file path to your JSON file.*

Skip to section **(4) Running BellaVista** for more information!

##### 3.3.3 Custom MERFISH

BellaVista can visualize datasets from custom MERFISH setups that were processed via the MERlin pipeline [2].

Standard MERlin output folder organization:

| | |
|---|---|
| %ANALYSIS_HOME% | Directory containing MERlin outputs |
| └ ExportBarcodes | Transcript coordinate locations and barcode ID |
| └ barcodes.csv |  |
| └ FiducialCorrelationWarp | Aligned FOV images |
| └ Images |  |
| └ aligned_images0.tif |  |
| └ … |  |
| └ aligned_images406.tif |  |
| └ RefineCellDatabases | HDF5 databases of cell/nuclear segmentation boundaries |
| └ features |  |
| └ feature_data_0.hdf5 |  |
| └ … |  |
| └ feature_data_406.hdf5 |  |
| └ … | |
| └ codebook.csv | Gene names/IDs for each barcode ID |
| └ microscope_parameters.json | Micron to pixel transformation |
| └ positions.csv | List of FOV microscope positions |

*Continued on next page* ⟶

From these outputs, it is possible to visualize tissue images, spatial transcript locations, and cell/nuclear segmentation boundaries.

To visualize tissue images, individual field-of-views (FOVs) must be stitched together. FOV images can be found in `FiducialCorrelationWarp/Images` in the MERlin output folder. Currently, BellaVista does not include a stitching pipeline. Stitching can be accomplished using image processing, utilizing Python packages such as NumPy [3] and Dask [4], or software such as BigStitcher [5]. The output stitched image must be a TIFF image, with individually stitched images for each channel you wish to visualize. The stitched images should be saved in the `%ANALYSIS_HOME%` directory.

Transcript locations and cell/nuclear segmentations exported by MERlin can be processed directly by BellaVista.

In order to visualize your MERFISH dataset in BellaVista, you will need to create a dataset-specific JSON configuration file containing paths to the MERlin outputs for your dataset. These output files will be processed to generate visualization files for BellaVista. Creating these visualization files will take a few minutes but only need to be created once. For subsequent runs, JSON parameter "create_inputs" can be set to `False`.

*Input file parameters on next page ⟶*

#### MERFISH-MERlin input file parameters

Below is a description of input file parameters for visualizing MERlin-processed MERFISH data. All input parameters are optional; if you are missing some input files, remove those input file parameters from the JSON file. BellaVista will skip the visualization of these features.

**Important:** All input file paths must be relative paths to *data_folder*.

- *data_folder* = the path to the `%ANALYSIS_HOME%` folder where the MERlin output files are stored.

---

**transcript_filename:** *string, default=None*
Relative path to CSV file containing decoded gene transcript spatial coordinates. If no file is provided, transcripts will not be processed or visualized.

**codebook:** *string, default=None*
Relative path to CSV containing map from "barcode_id" to gene ID. This is *required* to plot transcripts. If no file is provided, transcripts will not be processed or visualized.

**images:** *string or 1D array of strings, default=None*
Relative path to stitched image file(s). Must be a TIFF file. If no file is provided, images will not be processed or visualized.

- *When visualizing a single image, provide the file path as a string. For multiple images, pass them as a list of filenames. For example, use "DAPI.tif" for a single image or ["DAPI.tif", "PolyT.tif"] for multiple images.*

**microscope_parameters:** *string, default=None*
Relative path to JSON file containing microscope micron to pixel transform. This is *required* for displaying images. If no file is provided, no images can be visualized.

**positions_list:** *string, default=None*
Relative path to CSV file containing micron microscope positions for each FOV. This is *required* for displaying images. If no file is provided, no images can be visualized.

**z_plane:** *integer, default=0*
Z-plane of segmentations to plot. We suggest this match the z-plane of the image(s) you are visualizing. If no value is provided, the segmentations from the first z-plane will be processed and visualized.

**cell_segmentation:** *string, default=None*
Relative path to folder containing HDF5 cell segmentation boundaries. If no folder path provieded, no cell segmentations will be processed and visualized.

**nuclear_segmentation:** *string, default=0*
Relative path to folder containing HDF5 nuclear segmentation boundaries. If no folder path provieded, no nuclear segmentations will be processed and visualized.

*Continued on next page ⟶*

Certain features require multiple input files. Summary of required input files visualization for each feature:

**Transcripts**: "transcript_filename", "codebook"

**Images**: (1) "images" **&** (2) "microscope_parameters" **&** (3) "positions_list"

**Segmentations**: "cell_segmentation" and/or "nuclear_segmentation"

Example MERlin JSON configuration file:

```
{
    "system": "merlin",
    "data_folder": "/path/to/MERlin_ANALYSIS_HOME",
    "create_bellavista_inputs": true,

    "visualization_parameters": {
        "plot_image": true,
        "plot_transcripts": true,
        "plot_cell_seg": true
    },

    "input_files": {
        "transcript_filename": "ExportBarcodes/barcodes.csv",
        "codebook": "codebook.csv",
        "images": "polyT_z3.tif",
        "microscope_parameters": "microscope_parameters.json",
        "positions_list": "positions.csv",
        "z_plane": 3,
        "cell_segmentation": "RefineCellDatabases/features"
    }
}
```

Additional MERlin JSON configuration files and tutorials can be found on the documentation website: https://bellavista.readthedocs.io

#### Next steps:

Once your JSON is correctly configured for your dataset, you can run BellaVista in the terminal:

```
bellavista my_dataset.json
```

- *Replace "my_dataset.json" with the filename of the JSON you created. The JSON file argument should contain the file path to your JSON file.*

Skip to section **(4) Running BellaVista** for more information!

#### 3.4 Output Formats

The dataset input files to generate standardized data formats compatible with BellaVista. The standardized data are saved to a subfolder within the input data directory. Typically, the output data size is the same magnitude or smaller than the input data; however, we suggest having disk storage equivalent to the size of the input data available. For loading a typical-sized dataset, a computer with at least 16GB of memory is recommended.

Image files are converted to a pyramidal OME-Zarr next-generation file format (NGFF) utilizing compression of image chunks at multiple resolution levels for rapid loading and smooth transitioning between different magnifications as the user zooms in and out [6]. Gene transcript locations are stored in a dictionary mapping transcript coordinate locations to gene identities and segmentations are stored in an array. These arrays are stored in a pickle file format.

If an error occurs while processing any input file, BellaVista will stop the processing of this feature and record the error to `error_log.log` located in the `BellaVista_output` folder. Subsequently, continuing to process the next feature. BellaVista will automatically skip the visualization of any features that were not successfully processed.

*BellaVista_output folder organization on next page ⟶*

BellaVista_output subfolder organization:

| Path | Description |
| --- | --- |
| data_folder | Directory containing dataset input files |
| └ BellaVista_output | Subfolder containing BellaVista output formats |
| ├ OMEzarrimages | Multi-resolution pyramidal images |
| └ Images | |
| ├ s0 | |
| ├ s1 | |
| └ s2 | |
| ├ image_file_names.pkl | napari image layer names |
| ├ um_to_px_transforms.pkl | Micron-pixel transform + xy shifts for image alignment |
| ├ gene_dict.pkl | Transcript coordinates + napari gene layer names |
| ├ cell_boundary_coords.pkl | Cell segmentations |
| ├ nuclear_boundary_coords.pkl | Nuclear segmentations |
| ├ exceptions.json | Specifies which visualization features to skip (due to errors during processing input data) |
| └ error_log.log | Record of errors encountered during processing |

### 4 Running BellaVista

Once your JSON is correctly configured for your dataset, you can run BellaVista in the terminal:

```
bellavista /path/to/my_dataset.json
```

- *Replace "my_dataset.json" with the filename of the JSON you created. The JSON file argument should contain the file path to your JSON file.*

**Note**: It will take a few minutes to create the required data files.

The terminal will print updates & have progress bars for time consuming steps i.e.

```
Directory /Users/KosuriLab/xenium_mouse_brain_rep3/BellaVista_output
    does not exist -- creating the directory!
Creating BellaVista input files for 10x Genomics Xenium

Creating OME-Zarr image at 2024-12-09 21:10:46
Processing single image: morphology_mip.ome.tif
OME-Zarr image saved successfully at 2024-12-09 21:11:38

Calculating micron to pixel transform
Micron to pixel transform saved!

Creating transcripts
Processing genes: 100%|===========| 541/541 [00:04<00:00, 120.76it/s]

Creating cell boundaries
Processing cell boundaries: 100%|===========| 158047/158047 [00:12<00:00,
    12397.25it/s]
Cell boundaries saved!

No nuclear segmentation file provided, skipping segmentation processing.
    `plot_nuclear_seg` will remain false until a valid file is given.

BellaVista input files created!
```

*Continued on next page ⟶*

After the data has been successfully processed, BellaVista will then load the data into napari. Once successfully loaded, you should see the message `Data Loaded!` in the terminal.

A napari window should appear displaying the data similar to the image below:

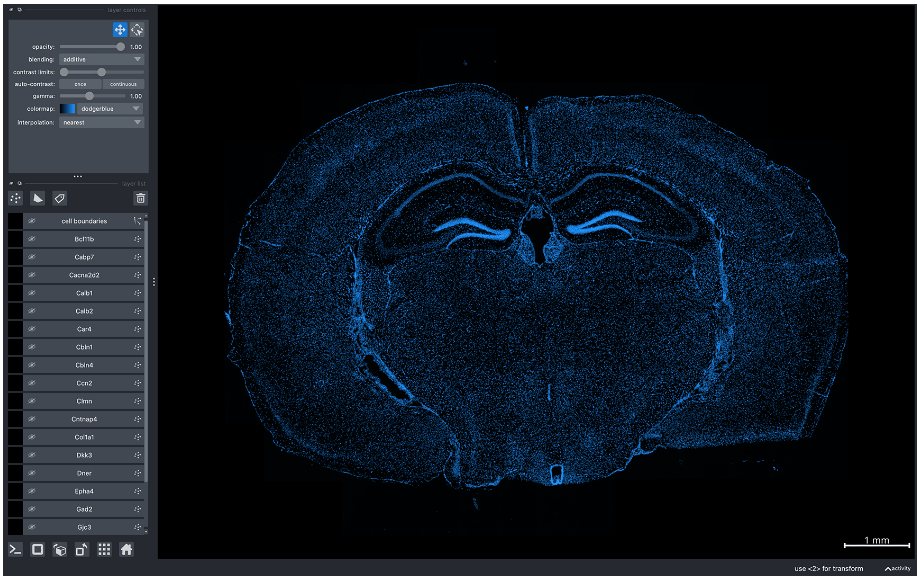

### 5 Generating Figures

Here you can find a list of commands that can be executed in the napari console. Additional commands and napari tips can be found in the FAQ on the documentation website: https://bellavista.readthedocs.io

#### 5.1 Using the napari console

To take screenshots and save current layer configurations, you can use the napari console. The napari console can be found in the bottom left of the GUI:

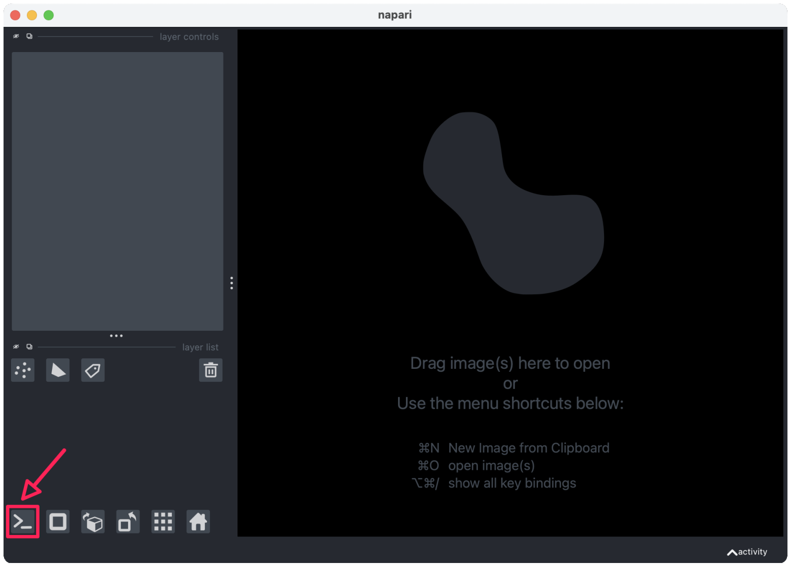

**Note:** To maximize content visible in the canvas, open the console in a separate window:

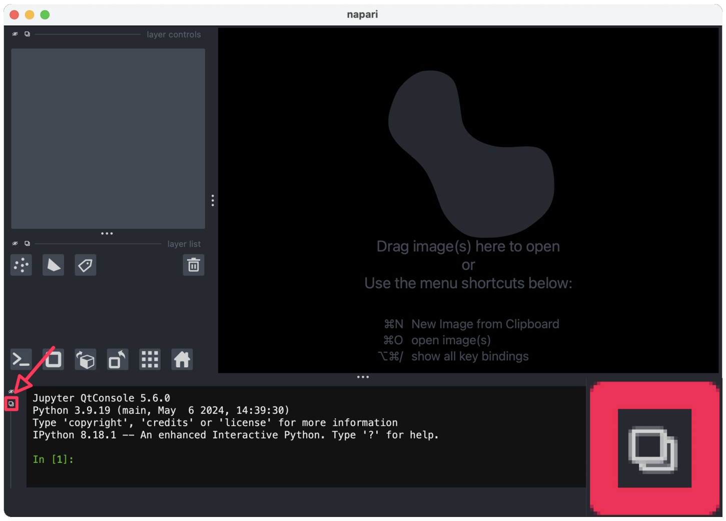

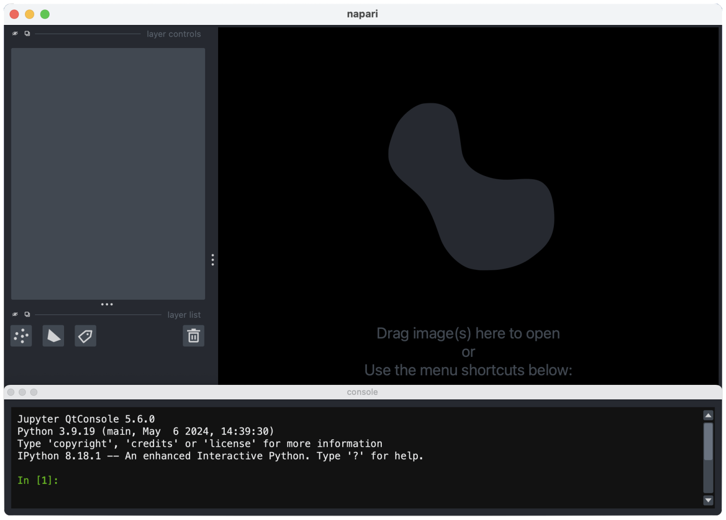

*Continued on next page* ⟶

#### 5.2 Useful Commands

Screenshot current canvas:

```
# save screenshot to folder
viewer.screenshot("/path/to/folder/napari_canvas.png")
```

Set saved screenshot resolution:

```
# set save resolution to 4K
save_resolution = (2160, 3840)

# save screenshot with 4K resolution to folder
viewer.screenshot("/path/to/folder/napari_canvas.png", size =
    save_resolution)
```

Save current camera position & zoom:

```
import pickle

# save current camera state
position_1 = {'center': viewer.camera.center, 'zoom': viewer.camera.zoom}
with open('/path/to/folder/position_1.pkl', 'wb') as f:
    pickle.dump(position_1, f)
```

Return to saved position:

```
import pickle

# load previous camera state
with open('/path/to/folder/position_1.pkl', 'rb') as f:
    position_1 = pickle.load(f)

# return to camera state from previous screenshot
viewer.camera.center, viewer.camera.zoom = position_1['center'],
    position_1['zoom']
```

Change transcript color & size:

```
import pickle

# change point color for "geneA" to magenta
viewer.layers["geneA"].face_color = "magenta"

# change color using hexcode
viewer.layers["geneA"].face_color = "#FF00FF"

# change point size for "geneA" to 2
viewer.layers["geneA"].size = 2
```

Save transcript colors for each gene:

```
import pickle

# store current transcript colors in a dictionary
gene_colors_dict = {}
gene_layers = [layer for layer in viewer.layers if isinstance(layer,
    napari.layers.Points)]
for gene_layer in gene_layers:
    gene_colors_dict[gene_layer.name] = gene_layer.face_color[0]

# save dictionary
with open('/path/to/folder/gene_colors_dict.pkl', 'wb') as f:
    pickle.dump(gene_colors_dict, f)
```

Load saved transcript colors:

```
import pickle

# load saved transcript colors
with open('/path/to/folder/gene_colors_dict.pkl', 'rb') as f:
    gene_colors_dict = pickle.load(f)

# assign gene transcript point colors
for gene in gene_colors_dict.keys():
    viewer.layers[gene].face_color = gene_colors_dict[gene]
```

Export point (gene) layers as svg file:

```
# save single layer as svg file
viewer.layers["geneA"].save("path/to/geneA.svg")

# save range of layers to single svg file
viewer.layers[5:10].save("path/to/5genes.svg")
```

**Important**: To export point layers to a SVG file, the point size must be an integer. If your point size is a float, you can change the point size using the commands in the example below.

Changing point sizes:

```
# check current point size for "geneA"
print(viewer.layers["geneA"].size)

# change point size for "geneA" to 1
viewer.layers["geneA"].size = 1
```

### 6 Additional Documentation

Further documentation and up-to-date information can be found on GitHub: https://github.com/pkosurilab/BellaVista

Additional example tutorials for each technology/platform, frequently asked questions (FAQ), and API information can be found on the documentation website: https://bellavista.readthedocs.io

The FAQ also includes a collection of helpful tips we've compiled based on user feedback. We will continue to update this list regularly. If you have suggestions that others might find helpful or if you're encountering issues not covered in the FAQ, please check the open issues or open a new issue in the GitHub repository: https://github.com/pkosurilab/BellaVista/issues
